# Mouse models of hereditary hemochromatosis do not develop early liver fibrosis in response to a high fat diet

**DOI:** 10.1101/319442

**Authors:** John Wagner, Carine Fillebeen, Tina Haliotis, Jeannie Mui, Hojatollah Vali, Kostas Pantopoulos

**Affiliations:** Lady Davis Institute for Medical Research, Jewish General Hospital, and Department of Medicine, McGill University, Montreal, Quebec, Canada; Department of Anatomy and Cell Biology, McGill University, Montreal, Quebec, Canada

**Keywords:** iron overload, hemochromatosis, NAFLD, NASH, liver fibrosis

## Abstract

Hepatic iron overload, a hallmark of hereditary hemochromatosis (HH), triggers progressive liver disease. There is also increasing evidence for a pathogenic role of iron in non-alcoholic fatty liver disease (NAFLD), which may progress to non-alcoholic steatohepatitis (NASH), fibrosis, cirrhosis and hepatocellular cancer. Mouse models of HH and NAFLD can be used to explore potential interactions between iron and lipid metabolic pathways. Hfe−/− mice, a model of moderate iron overload, were reported to develop early liver fibrosis in response to a high fat diet. However, this was not the case with Hjv−/− mice, a model of severe iron overload. These data raised the possibility that the *Hfe* gene may protect against liver injury independently of its iron regulatory function. Herein, we addressed this hypothesis in a comparative study utilizing wild type, Hfe−/−, Hjv−/− and double Hfe−/−Hjv−/− mice. The animals, all in C57/BL6 background, were fed with a high fat diet for 14 weeks and developed hepatic steatosis, associated with mild iron overload. Hfe co-ablation did not sensitize steatotic Hjv-deficient mice to liver injury. Moreover, we did not observe any signs of liver inflammation or fibrosis even in single steatotic Hfe−/− mice. Ultrastructural studies revealed a reduced lipid and glycogen content in Hjv−/− hepatocytes, indicative of a metabolic defect. Interestingly, glycogen levels were restored in double Hfe−/−Hjv−/− mice, which is consistent with a metabolic function of Hfe. We conclude that hepatocellular iron excess does not aggravate diet-induced steatosis to steatohepatitis or early liver fibrosis in mouse models of HH, irrespectively of the presence or lack of Hfe.

## Introduction

Non-alcoholic fatty liver disease (NAFLD) represents the hepatic component of the metabolic syndrome (type 2 diabetes, obesity, hyperlipidemia, hypertension), and constitutes the most frequent liver disease in Western countries [1, 2]. NAFLD is characterized by excessive fat accumulation in hepatocytes in the absence of other causes of liver disease, such as alcohol abuse or viral hepatitis. In approximately 30% of patients, NAFLD progresses from simple steatosis to non-alcoholic steatohepatitis (NASH), a chronic inflammatory condition that may further lead to liver fibrosis, cirrhosis and hepatocellular carcinoma (HCC). NAFLD patients often exhibit perturbed iron metabolism and accumulate liver iron deposits, which increases the risk for liver fibrosis [3, 4]. Consequently, manipulation of iron metabolic pathways may offer a promising therapeutic target for NAFLD [5].

Systemic iron balance is controlled by hepcidin, a liver-derived iron regulatory hormone [6]. Hepcidin limits iron efflux to the bloodstream by binding to the iron exporter ferroportin in tissue macrophages, intestinal enterocytes and other target cells, which leads to ferroportin internalization and degradation. The expression of hepcidin is induced in response to iron stores, inflammatory signals and other stimuli. Iron regulation of hepcidin involves bone morphogenetic proteins (BMPs) and the SMAD signaling cascade [7]. Genetic defects in the hepcidin pathway underlie the development of hereditary hemochromatosis, an endocrine disorder of systemic iron overload that is caused by loss of feedback regulation in iron absorption and systemic iron traffic [8, 9]. The most common form of hemochromatosis is associated with mutations in the HFE, an atypical major histocompatibility class I molecule. Inactivation of the HJV (hemojuvelin), a BMP co-receptor, leads to early onset juvenile hemochromatosis, characterized by more severe iron overload. Both HFE and HJV operate as upstream regulators of iron signaling to hepcidin [10].

Hemochromatosis patients exhibit excessive iron accumulation within hepatocytes, which predisposes them to liver fibrosis and progression to end stage liver disease [11]. Hfe−/− and Hjv−/− mice recapitulate the relatively milder or severe iron overload of patients with adult or juvenile hemochromatosis, respectively, but do not develop spontaneous early liver disease. Interestingly, Hfe- /- mice manifested a NASH-like phenotype and early liver fibrosis after feeding a high fat diet for 8 weeks, which was only partially attributed to iron [12]. On the other hand, a 12-week high fat diet intake caused liver steatosis but did not promote steatohepatitis or liver fibrosis in Hjv−/− mice, in spite of severe hepatocellular iron overload [13]. These data raised the possibility for a protective function of Hfe against metabolic liver disease. Herein, we explored this hypothesis by comparing pathophysiological responses of mice with single or combined Hfe/Hjv deficiency to high fat diet.

## Materials and Methods

### Animals

Four-week old male wild-type, Hfe−/−, Hjv−/− and double Hfe−/−Hjv−/− mice [14] (n=10 for each genotype) were fed after weaning with a standard diet (Harlad Teklad 2018) or a high fat diet (Harlad Teklad TD.88137) for 14 weeks. A group of 10-week old male wild type mice were treated for 6 weeks with CCl_4_ to develop liver fibrosis [15]. All animals were on C57BL/6 genetic background. They were housed in a temperature-controlled environment (22 ± 1° C, 60 ± 5% humidity), with a 12-hour light/dark cycle and were allowed *ad libitum* access to diets and drinking water. At the endpoint, the mice were sacrificed by CO_2_ inhalation. Blood was obtained by cardiac puncture and clotted at room temperature for 1 h. Serum was separated by centrifugation (2000 g for 10 min), snap frozen in liquid nitrogen and stored at -80°C for biochemical analysis. Livers were rapidly excised and tissue sections were snap frozen in liquid nitrogen and stored at -80°C for biochemical studies. Other liver sections were fixed in 10% neutral-buffered formalin and embedded in paraffin for histological and ultrastructural studies. All animal procedures were approved by the Animal Care Committee of McGill University (protocol 4966).

### Serum biochemistry

Iron, transferrin saturation, total iron binding capacity (TIBC), ferritin, glucose, triglycerides, cholesterol, HDL cholesterol, alanine transaminase (ALT) and aspartate transaminase (AST) were measured with a Roche Hitachi 917 Chemistry Analyzer at the Biochemistry Department of the Jewish General Hospital.

### Liver biochemistry

Liver extracts were analyzed for non-heme iron by the ferrozine assay [16], and for the presence of collagen by a colorimetric hydroxyproline assay (QuickZyme Biosciences), according to the manufacturer’s recommendations.

### Histopathology and immunohistochemistry

Deparaffinized liver sections were stained with hematoxylin and eosin (H&E), Perls’ Prussian blue or Masson’s trichrome to assess tissue architecture, iron deposits or collagen, respectively. Expression of α-smooth muscle actin (α-SMA) was analyzed by immunohistochemistry, as previously described [15].

### Transmission electron microscopy

Liver sections were prepared for ultrastructural studies and analyzed by transmission electron microscopy (TEM) as described [13].

### Statistics

Statistical analysis was performed with the GraphPad Prism software (v. 5.0d). Quantitative data are expressed as mean ± standard error of the mean (SEM). Statistical analysis across multiple groups (genotypes and diets) was performed by two-way ANOVA with Bonferroni post-test comparison. A probability value p<0.05 was considered statistically significant.

## Results

### Responses of mouse models of hemochromatosis to a high fat diet

Hfe−/− mice represent a model of the most common and relatively milder form of hemochromatosis, while Hjv−/− and double Hfe−/−Hjv−/− mice develop more severe iron overload [14]. Thus, when compared to isogenic wild type control animals, serum iron, ferritin and transferrin saturation were modestly elevated in Hfe−/− and substantially augmented in Hjv−/− and double Hfe−/−Hjv−/− mice on a standard diet (Fig. 1B-D). Feeding a high fat diet (HFD) immediately after weaning for 14 weeks resulted an approximately 35% body weight gain in all mice irrespective of genotype, as compared to age- and sex-matched controls fed a standard diet (Fig. 1A). HFD intake was associated with slight increases in serum iron (Fig. 1B) and TIBC (Fig. 1E) and commensurate drops in ferritin (Fig. 1C) and transferrin saturation (Fig. 1D).

**Fig. 1.**
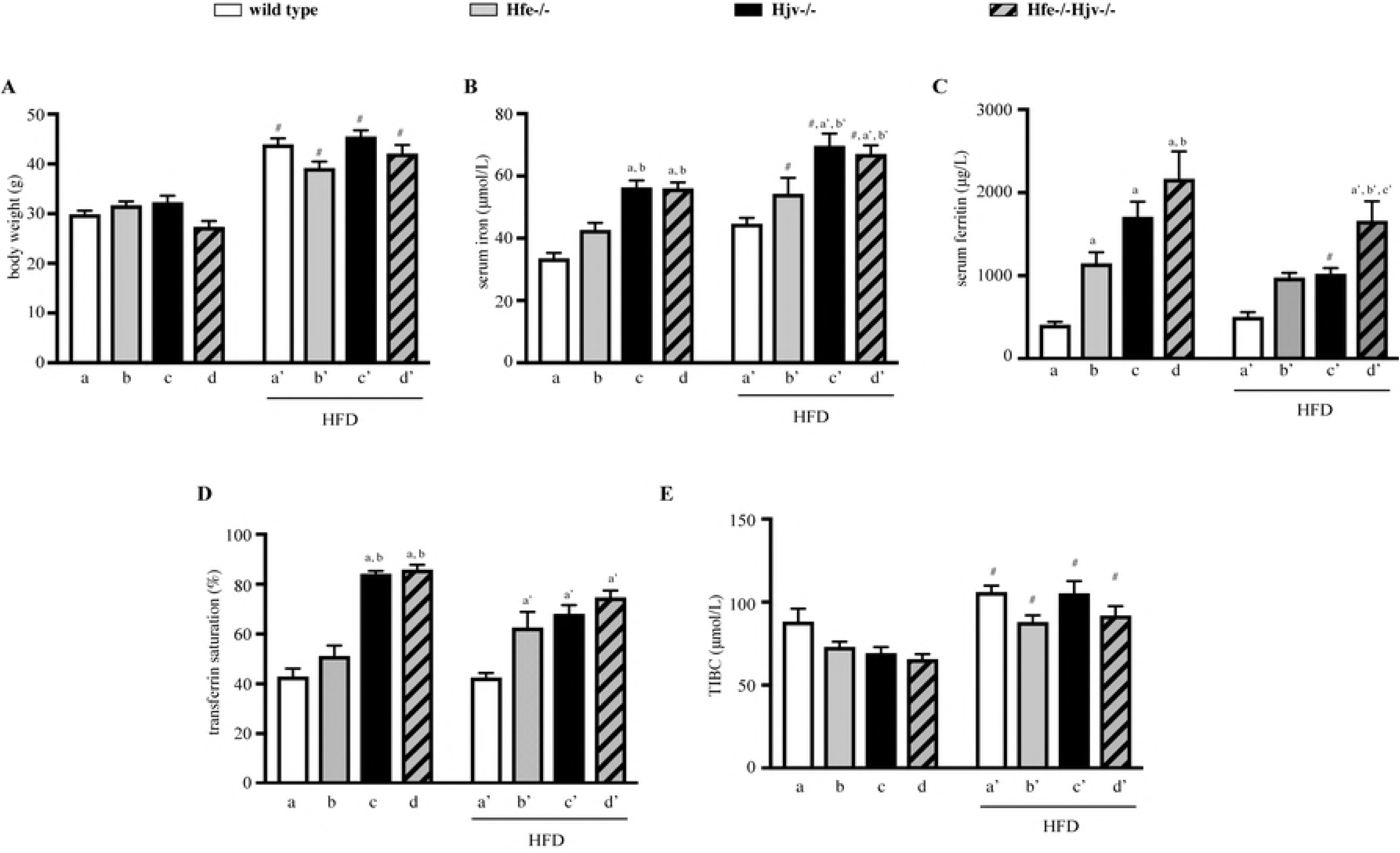
Effects of high fat diet on body weight and serum iron parameters. Wild type (wt), Hfe−/−, Hjv−/− and double Hfe−/−Hjv−/− mice (n=10 per group, all males in C57BL/6 genetic background) were placed immediately after weaning on a standard diet, or a high fat diet (HFD). After 14 weeks, body weights were monitored, the animals were sacrificed, and sera and liver tissues were obtained for analysis. (A) Body weights of the mice; (B) serum iron; (C) serum ferritin; (D) transferrin saturation; and (E) total iron binding capacity (TIBC). All data are presented as the mean ± SEM. Statistical analysis was performed by two-way ANOVA. Statistically significant differences (p<0.05) across genotypes (versus columns a, b, a’, b’, c’) are indicated by a, b, a’, b’, c’ and across diets by #.

In addition, HFD intake promoted an increase in levels of serum glucose, cholesterol and HDL-cholesterol, but not triglycerides, in all genotypes (Fig. 2A-D). The effects on cholesterol and HDL-cholesterol appeared less pronounced in Hfe−/− mice. These animals were also spared from HFD-mediated increases in serum transaminases ALT and AST, which were evident in wild type, Hjv−/− and Hfe−/−Hjv−/− mice (Fig. 2E-F).

**Fig. 2.**
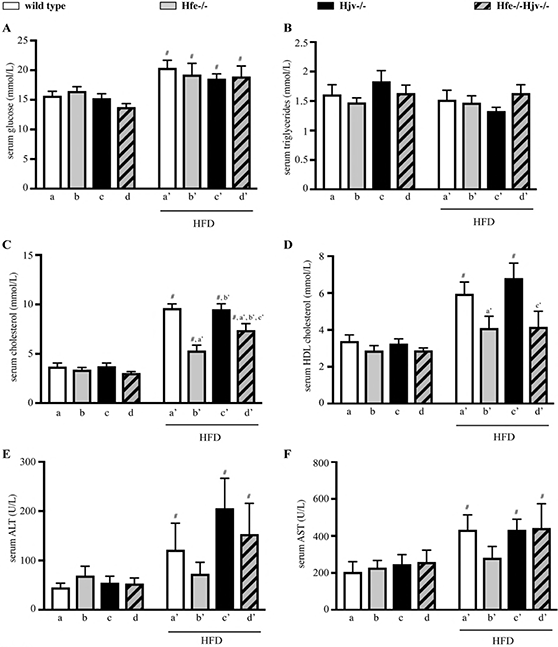
Effects of high fat diet on serum biochemistry. Sera from mice described in Fig. 1 were analysed for: (A) glucose; (B) triglycerides; (C) cholesterol; (D) HDL-cholesterol; (E) ALT; and (F) AST. All data are presented as the mean ± SEM. Statistical analysis was performed by two-way ANOVA. Statistically significant differences (p<0.05) across genotypes (versus columns a’, b’, c’) are indicated by a’, b’, c’ and across diets by #.

Livers of all mice on HFD were enlarged by ~40% (data not shown) and exhibited microvesicular steatosis (Fig. 3). There were no differences among genotypes in the degree and the pattern of fat deposition. Histological analysis did not reveal any large inflammatory foci or clusters of immune cells in any of the examined livers. The above data suggest that the high fat diet promotes similar obesity and liver steatosis in wild type mice and mouse models of hemochromatosis.

**Fig. 3.**
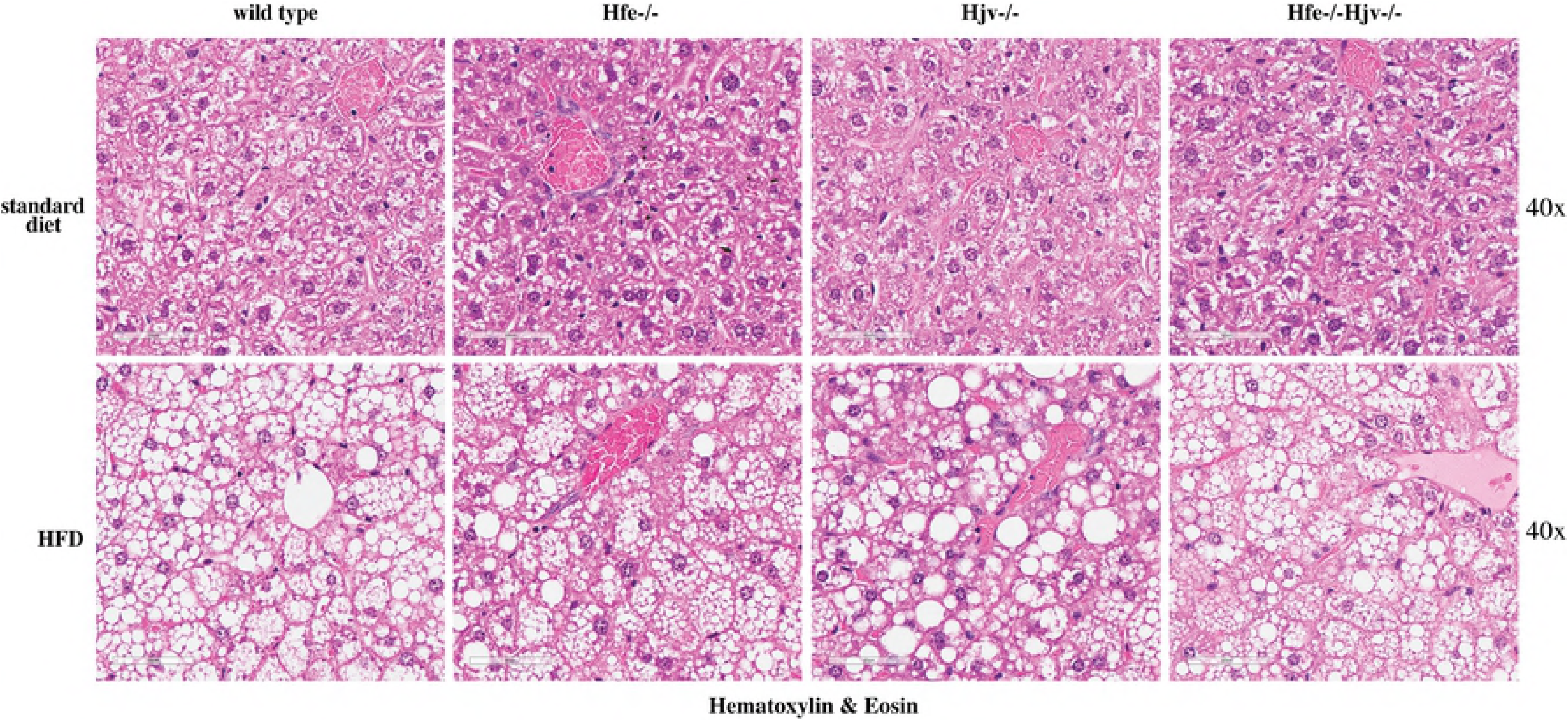
High fat diet promotes liver steatosis without necroinflammation in mouse models of hemochromatosis. Liver sections from the mice described in Fig. 1 were stained with H&E. Original magnification: 40x.

### Does hepatic iron overload trigger progression of steatosis to liver fibrosis?

As expected, mouse models of hemochromatosis developed hepatic iron overload (Fig. 4). This was relatively mild in Hfe−/− mice and more intense in Hjv−/− and double Hfe−/−Hjv−/− counterparts, in agreement with earlier data [14]. Liver iron accumulation was reduced in all animals fed the HFD, as previously observed [12, 13, 17, 18], and this effect was more pronounced in Hjv−/− and double Hfe−/−Hjv−/− mice (Fig. 4A). Notably, HFD-fed Hfe−/−, Hjv−/− and Hfe−/−Hjv−/− mice had similar liver iron content. This remained elevated compared to control HFD-fed wild type mice. Iron deposits were visualized by Perls’ Prussian blue staining in hepatocytes of HFD-fed mouse models of hemochromatosis but not wild type controls (Fig. 4B).

**Fig. 4.**
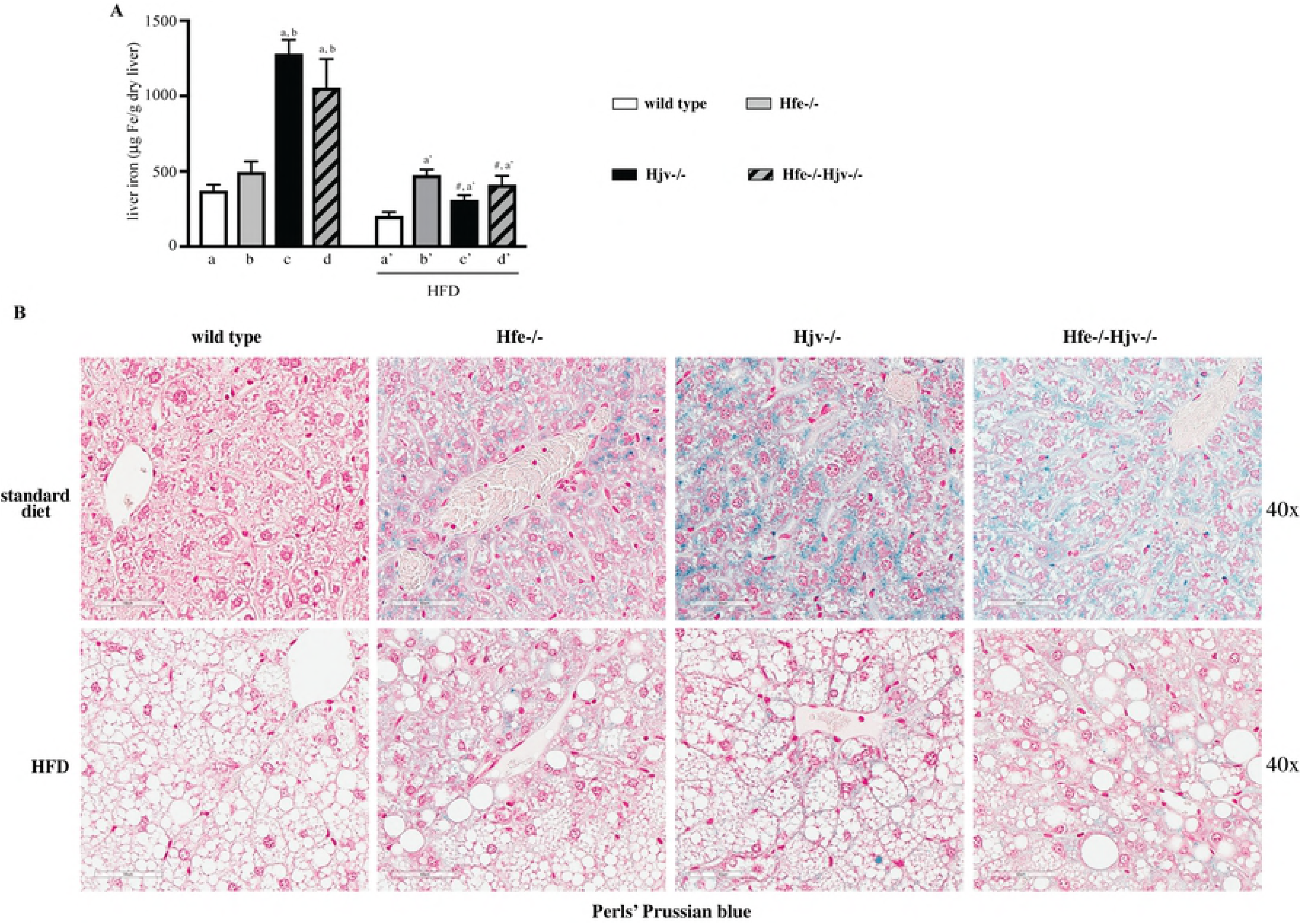
Evaluation of hepatic iron content in mouse models of hemochromatosis on standard or high fat diet. Livers from the mice described in Fig. 1 were used to quantify and histologically assess iron content. (A) Quantification of non-heme iron by the ferrozine assay. Data are presented as the mean ± SEM. Statistical analysis was performed by two-way ANOVA. Statistically significant differences (p<0.05) across genotypes (versus columns a, b, a’) are indicated by a, b, a’ and across diets by #. (B) Histological detection of iron deposits by staining with Perls’ Prussian blue. Original magnification: 40x.

Hepatocellular iron overload was not associated with liver fibrosis in any of the mice fed standard diet or HFD. The presence of collagen was assessed histologically by Masson’s trichrome staining (Fig. 5A) and biochemically by the hydroxyproline assay (Fig. 5B). Liver samples from CCl_4_-treated wild type mice served as positive control for liver fibrosis. Immunohistochemical analysis showed expression of α-SMA, a marker of activated hepatic stellate cells, in livers of Hjv−/−, Hfe−/− Hjv−/− and to a smaller extent also Hfe−/− mice (Fig. 6). In addition, HFD intake promoted α-SMA expression in livers of wild type mice. Thus, either hepatic iron overload or steatosis appear to activate fibrogenic responses in mouse models of hemochromatosis; however, their combination does not dramatically accelerate them to promote rapid progression of liver disease.

**Fig. 5.**
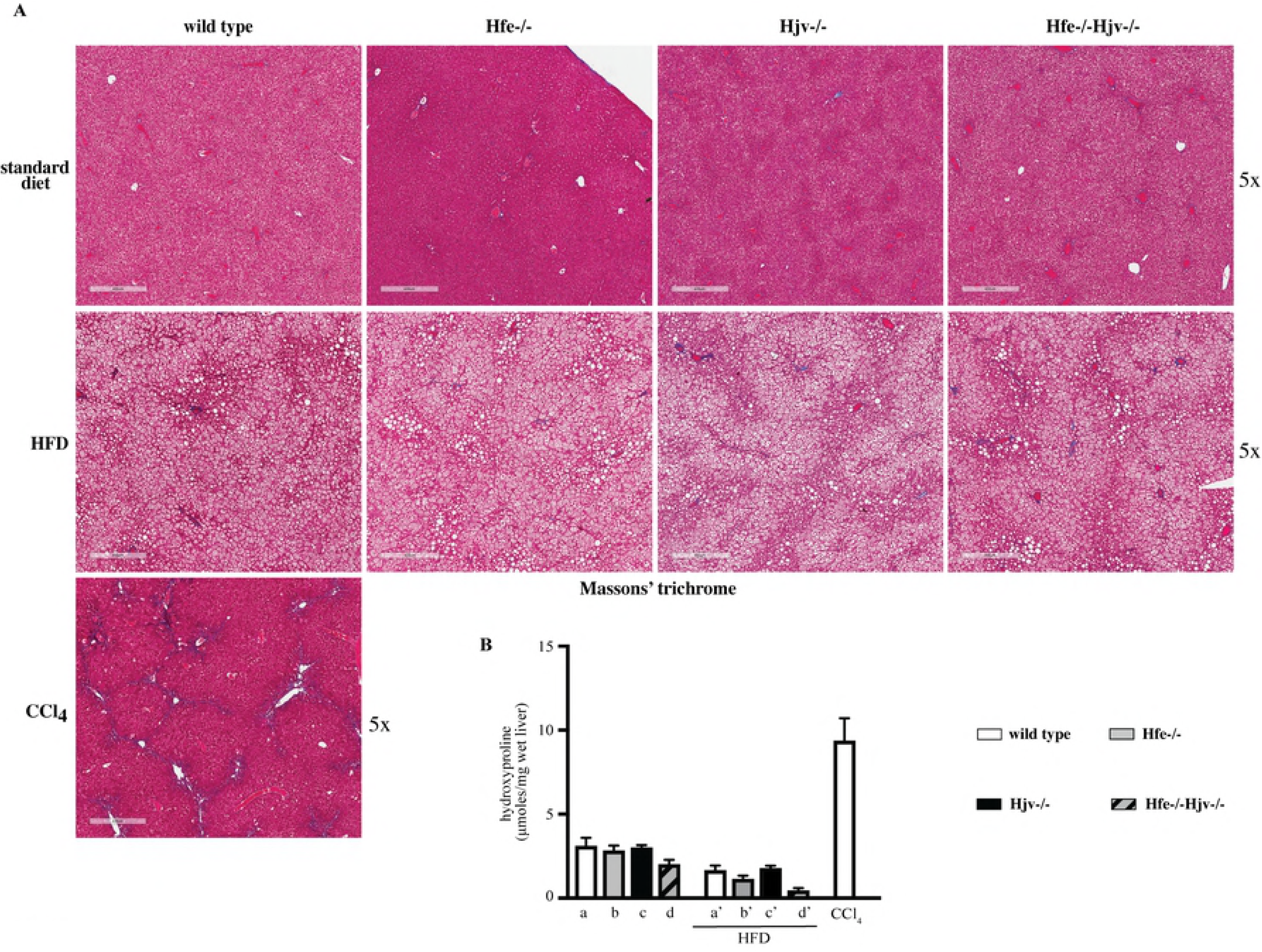
High fat diet does not promote liver fibrosis in mouse models of hemochromatosis. Livers from the mice described in Fig. 1 were used to histologically assess fibrosis and to quantify collagen. Livers from mice treated with CCl_4_ were used as positive control. (A) Staining with Masson’s trichrome. Original magnification: 5x. (B) Quantification of collagen by the hydroxyproline assay. Data are presented as the mean ± SEM. Statistical analysis was performed by two-way ANOVA.

### Ultrastructural studies

Analysis of the liver tissue by TEM corroborated the absence of inflammation or fibrosis in hemochromatotic and steatotic mice. Data obtained from animals on standard diet are shown in Fig. 7 (top). Wild type hepatocytes exhibited a normal architecture and contained regular lipid droplets and glycogen granules. On the other hand, some mitochondria displayed darker contrast suggesting the possible presence of lipids stained by osmium tetroxide. The same feature was observed in Hfe−/− hepatocytes, but some mitochondria were swollen (arrow). Hjv−/− hepatocytes displayed well-preserved mitochondria with much less dense glycogen and reduced lipid droplets. ER appears to be less organized and located more around mitochondria. Interestingly, in Hfe- /-Hjv−/− hepatocytes the reduced lipid content persisted but glycogen density and mitochondrial appearance were normalized.

Data from mice on HFD are depicted in Fig. 7 (bottom). The number and size of lipid droplets was increased in all genotypes. Nevertheless, the number of fat vesicles was relatively higher in Hfe−/−, and the overall fat content lower in Hjv−/− hepatocytes. The appearance of mitochondria was improved in Hfe−/− hepatocytes but the glycogen density was reduced compared to that from mice on standard diet. By contrast, some mitochondria looked abnormal in Hjv−/− hepatocytes (arrows), the glycogen density remained low, and the ER got disorganized in response to the HFD. With respect to mitochondria and ER, hepatocytes from Hfe−/−Hjv−/− and Hfe−/− mice on HFD had a similar phenotype. Glycogen density appears reduced. Taken together, the transmission electron microscopy data reveal distinct ultrastructural features among mouse models of hemochromatosis and raise the possibility for metabolic functions of Hfe and Hjv.

## Discussion

Excessive accumulation of iron or fat in hepatocytes are known pathogenic factors for liver injury. Clinical data suggested a synergism of hepatocellular iron overload and steatosis in liver disease progression to liver fibrosis [19–21]. We sought to validate these findings using Hjv−/− mice, a model of severe, early onset juvenile hemochromatosis. However, feeding Hjv−/− mice with a HFD did not cause progression of liver steatosis to steatohepatitis or fibrosis, in spite of profound iron overload [13]. In another study, HFD intake triggered early liver fibrosis in Hfe−/− mice, a model of relatively mild, late onset adult hemochromatosis [12]. Hfe−/− mice also manifested exacerbated hepatotoxicity following combined HFD and alcohol intake [22].

Based on these findings, we reasoned that Hfe may exert a potential iron-independent function in liver disease progression and utilized herein double Hfe−/−Hjv−/− mice to explore this hypothesis. Hfe−/−Hjv−/− and Hjv−/− mice exhibit an indistinguishable iron overload phenotype, suggesting that the *Hjv* gene is epistatic to *Hfe* in iron homeostasis [14]. Thus, comparing the responses of these animals to a HFD would provide insights on the role of Hfe in metabolic liver disease. To this end, isogenic sex and age-matched wild type control, Hfe−/−, Hjv−/− and double Hfe−/−Hjv−/− mice were exposed to a HFD for 14 weeks. HFD intake triggered obesity and liver steatosis in all genotypes but did not promote steatohepatitis or liver fibrosis in any of them (Figs. 3 and 5). These data do not support an iron-independent protective role of Hfe against liver disease progression.

The Hfe−/− mice used in this work and in [12] shared the same genetic background (C57BL/6). In our experiments, exposure to the HFD started a bit earlier (immediately after weaning vs at the 6th week of age) and lasted considerably longer (14 vs 8 weeks); this setting is expected to be more favorable to liver injury. Thus, we speculate that the discordant results reported herein and in [12] may be related to small variations in the HFD content. It will be interesting to examine responses of Hfe−/− Hjv−/− and Hjv−/− mice to the HFD used in [12]. The data with Hjv−/− mice are largely consistent with our previous findings [13]. However, we did not observe reduced body weight gain of Hjv−/− compared to wild type mice in response to HFD (Fig. 1A), contrary to data in [13]. Presumably, this discrepancy is related to the different timing the mice were placed on HFD (immediately after weaning vs at the 10^th^ week of age) and the duration of HFD feeding (14 vs 12 weeks).

Our data suggest that hepatocellular iron overload does not aggravate liver steatosis to steatohepatitis or fibrosis in the mouse models of hemochromatosis, at least within the time frame of 14 weeks. It is conceivable that iron’s hepatotoxicity is manifested at later time points. It should be noted that either iron or fat accumulation trigger fibrogenic responses on their own right, as illustrated by the induction of α-SMA (Fig. 6), a marker of hepatic stellate cell activation to collagen-secreting myofibroblasts [23]. However, their combination does not appear to aggravate or accelerate these responses in mice.

**Fig. 6.**
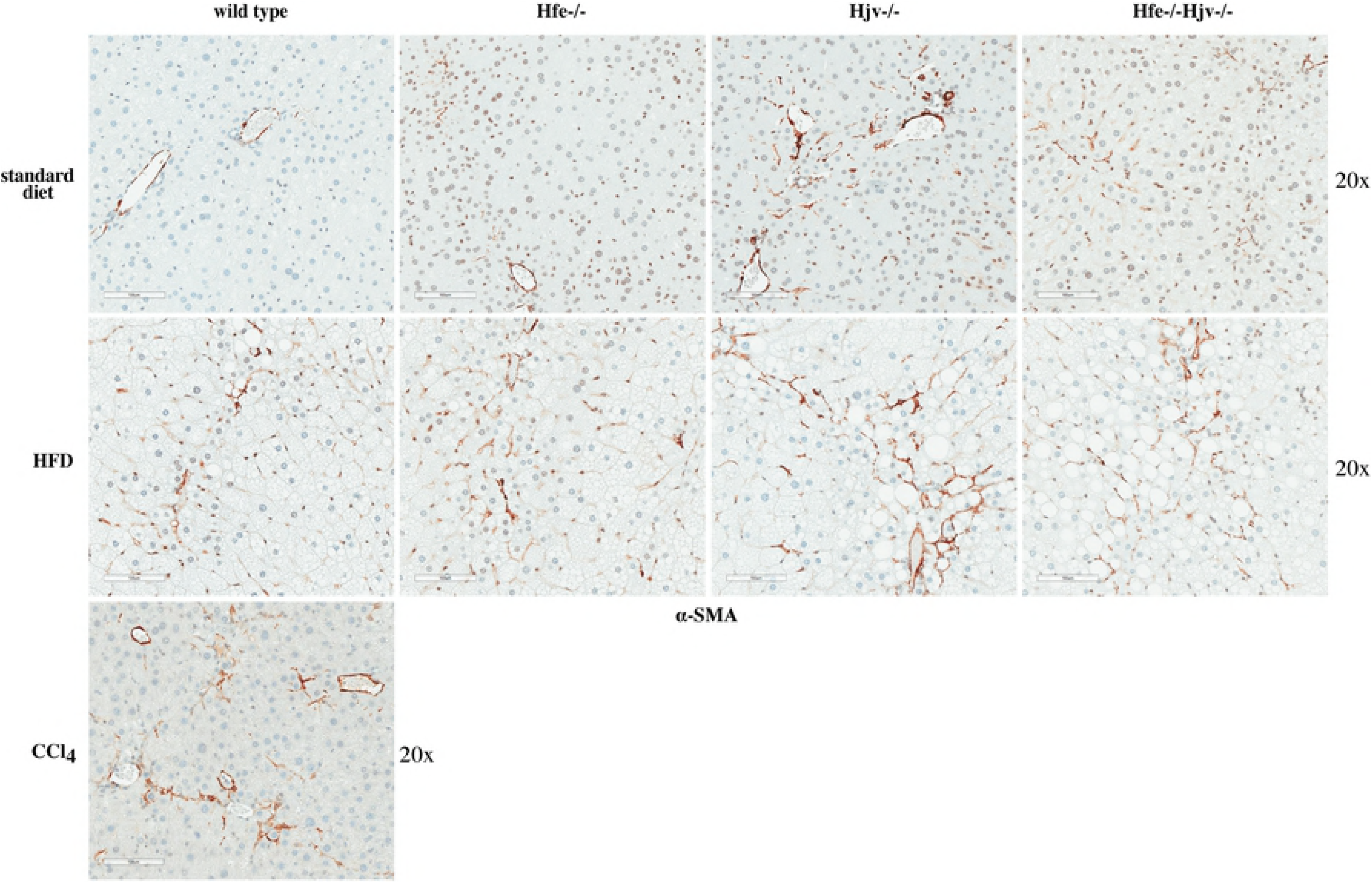
Hepatic iron overload or steatosis trigger activation of hepatic stellate cells. Liver sections from the mice described in Figures 1 and 5 were used for immunohistochemical detection of a smooth muscle actin (a-SMA), a marker of hepatic stellate cell activation. Original magnification: 20x.

Another issue that warrants consideration is that iron overload was not excessive in the HFD-fed mice, as compared to control animals of the same genotype on standard diet (Fig. 4). This is consistent with the reported reduced iron absorption/accumulation in mice fed high fat diets [12, 13, 17, 18], but is also related to the fact that the standard diet used herein had a ~4-fold higher iron content than the HFD (200 vs 50 ppm). Nevertheless, all HFD-fed hemochromatotic mice of our study had pathological hepatic iron content, which is evident by the positive Perls’ staining (Fig. 4B). Considering the ~40% increased size of the steatotic livers, the quantitative ferrozine assay as expressed in μg of iron per gram of dry liver (Fig. 4A) probably underestimates the true hepatocellular iron concentration. In any case, the low degree of iron overload is a limitation of this study and could at least partly explain the lack of iron pathogenicity, and also the discrepancy with the previous findings in Hfe−/− mice [12]. Thus, it will be important to evaluate a potential dose-dependent synergy of iron overload in liver disease progression in the context of the steatotic liver. It should, however be noted that supplementation of the HFD with 2% carbonyl iron did not promote liver fibrosis in Hjv−/− mice [13].

Assuming that even a low degree of hepatocellular iron overload is often pathogenic in humans, our results do not recapitulate clinical data on NAFLD in hemochromatosis patients [19–21]. This notion underlines known shortcomings of mouse models. Thus, it is well established that genetic mouse models of hemochromatosis do not develop spontaneous liver complications due to hepatocellular overload, contrary to hemochromatosis patients. On the other hand, dietary iron overload was recently shown to promote steatohepatitis in genetically obese Lepr^db/db^ mice; notably excess iron accumulated predominantly in reticuloendothelial cells [24]. This result is in line with clinical studies establishing a pathogenic role of reticuloendothelial iron overload, which has been documented in many NAFLD patients [25–27].

On a final note, the TEM analysis in Fig. 7 uncovers marked differences in hepatocyte ultrastructure of Hfe−/− and Hjv−/− mice. It identified apparent mitochondrial abnormalities in Hfe−/− hepatocytes, and confirmed the previously reported [13] reduction in fat and glycogen stores in Hjv−/− hepatocytes. Strikingly, glycogen density and mitochondrial morphology were corrected in Hfe−/−Hjv- /- hepatocytes. These findings are consistent with metabolic functions of Hfe and Hjv, most likely unrelated to their iron regulatory activities. Future work is expected to shed more light on relevant biochemical mechanisms and pathophysiological implications.

**Fig. 7.**
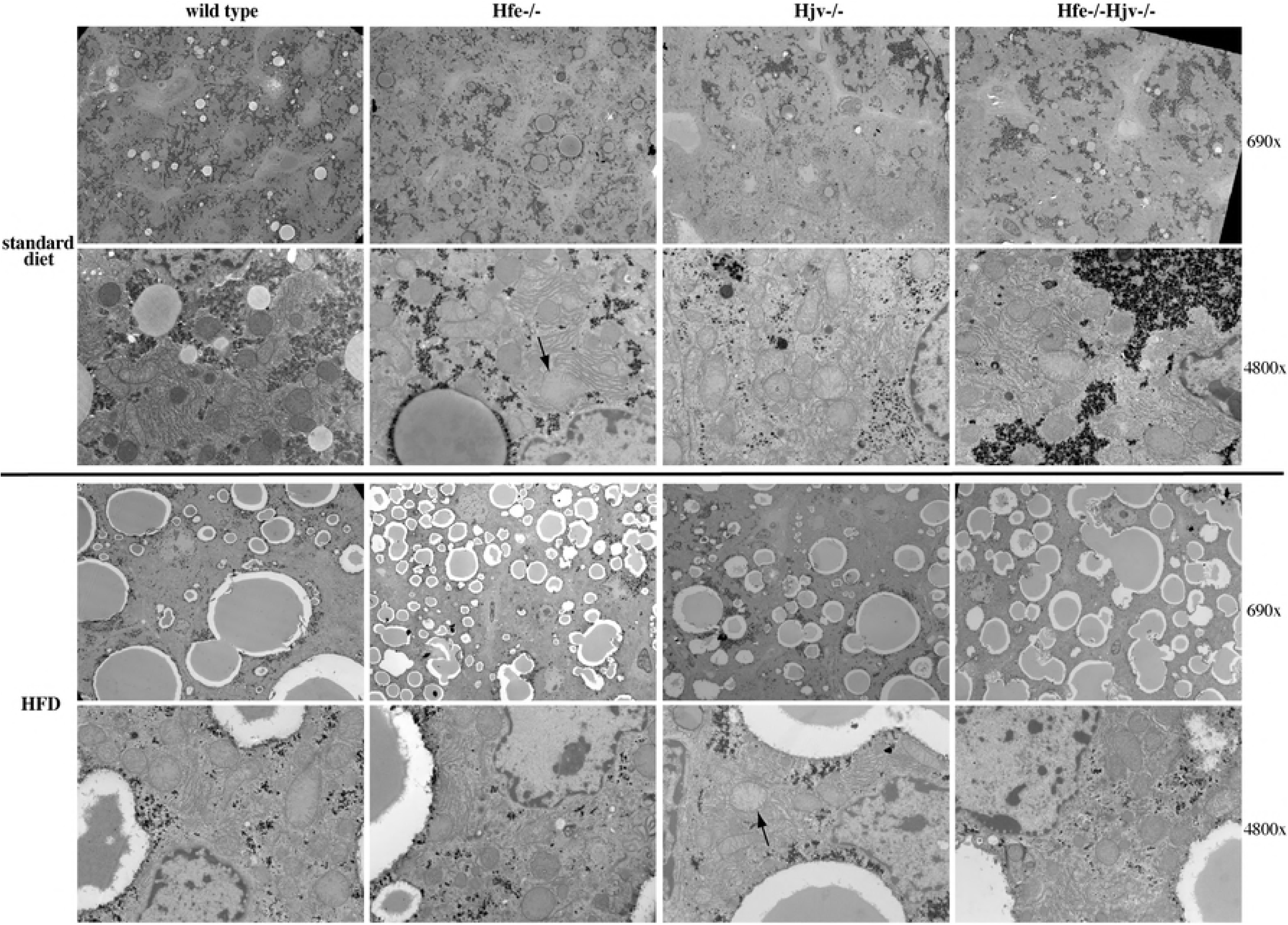
Ablation of Hfe or Hjv is associated with ultrastructural alterations in hepatocytes. Liver sections from the mice described in Fig. 1 were analyzed by transmission electron microscopy (TEM). Two magnifications (690x and 4800x) are shown for each sample. Arrows indicate mitochondria with abnormal morphology.

## Acknowledgements

We thank Drs. Naciba Benlimame for assistance with histology. This work was supported by a grant from the Canadian Institutes for Health Research (CIHR; MOP-86514).

## Author contributions

JW, CF and JM performed research and analyzed data; TH and HV analyzed data; KP designed and supervised research and wrote the paper.

